# Chromatin Assembly Factor 1 is required for normal structure and function of facultative heterochromatin in *Neurospora crassa*

**DOI:** 10.64898/2026.06.12.731976

**Authors:** Eduardo V. Torres, Rochelle E. Yap, Aileen R. Ferraro, Collin D. Link, Jacqueline F. Pelham, Zachary A. Lewis

## Abstract

Polycomb Repressive Complex 2 (PRC2) is a conserved epigenetic regulator that represses gene expression through methylation of histone H3 lysine 27 (H3K27me3). In animals, plants, and some fungi, PRC2-directed facultative heterochromatin plays essential roles in development and cellular differentiation. Here, we show that the replication-dependent histone chaperone Chromatin Assembly Factor 1 (CAF-1) is required for proper structure and function of facultative heterochromatin in the model fungus *Neurospora crassa*. Loss of CAF-1 causes widespread transcriptional misregulation, particularly within PRC2-repressed regions, and leads to redistribution of H3K27me3, reduced ASH1-dependent H3K36 methylation, and accumulation of chromatin marks associated with active transcription. CAF-1 was not required for repressive histone methylation within constitutive heterochromatin. A double mutant lacking both CAF-1 and PRC2 components displayed a synergistic silencing defect, suggesting these complexes make distinct contributions to facultative heterochromatin. Together, our findings indicate that CAF-1 works in concert with PRC2 to silence transcription within *N. crassa* facultative heterochromatin domains.

## Introduction

Epigenetic mechanisms such as histone post-translational modifications (PTMs) play central roles in eukaryotic gene regulation (Allis and Jenuwein 2016). Histone PTMs control gene expression by modulating DNA interactions with transcription factors, RNA polymerases, and other chromatin active enzymes or by directing the assembly of specialized chromatin domains that inhibit transcription (Allis and Jenuwein 2016). Heterochromatin refers to specialized chromatin states that are enriched for repressive histone PTMs and are transcriptionally inert (Trojer and Reinberg 2007; Saksouk et al. 2015; Allshire and Madhani 2018; Janssen et al. 2018; Grewal 2023). Heterochromatin can be further subdivided into constitutive heterochromatin, which is typically associated with repeat-rich, gene-poor regions of the genome, or facultative heterochromatin, which is gene-rich and dynamically repressed. Within facultative heterochromatin, genes are tightly repressed under most conditions but can be expressed in a cell type- or context-specific manner (Trojer and Reinberg 2007; Akilli et al. 2024; Rosas et al. 2026).

In many organisms, facultative heterochromatin is established and maintained by Polycomb Group (PcG) proteins, which assemble into multi-subunit chromatin complexes that deposit repressive histone modifications (Grossniklaus and Paro 2014; Schuettengruber et al. 2017; Tamburri et al. 2024). Polycomb Repressive Complex 2 (PRC2) catalyzes histone H3 lysine 27 trimethylation (H3K27me3), and PRC1 ubiquitylates H2A (Blackledge and Klose 2021; Tamburri et al. 2024). In plants and animals, PRC-dependent gene repression controls important processes including multicellular development, X-chromosome inactivation, and vernalization (Grossniklaus and Paro 2014; Zylicz and Heard 2020). Although PRC1 was apparently lost in an early fungal ancestor, PRC2 and H3K27me3 are present in diverse fungal lineages, with the model yeasts *Saccharomyces cerevisiae* and *Schizosaccharomyces pombe* representing notable exceptions (Erlendson et al. 2017). In fungi that have retained PRC2, H3K27me3 is linked to repression of genes associated with sexual development, secondary metabolism, and plant pathogenesis (Connolly et al. 2013; Collemare and Seidl 2019; Ridenour et al. 2020; Zhang et al. 2021; Fraser and Whitehall 2022; Deaven et al. 2025; Janevska et al. 2026). Despite the central role of PRC2 in eukaryotic gene regulation, the genes and mechanisms that establish and maintain facultative heterochromatin remain incompletely understood.

The filamentous fungus *Neurospora crassa* has emerged as a powerful model for dissecting the mechanisms that establish and maintain facultative heterochromatin (Lewis 2017). H3K27me3 covers ∼7% of the *N. crassa* genome and is enriched in large, multi-gene domains that are biased toward chromosome ends (Jamieson et al. 2013). Genetic studies have identified multiple factors required for normal facultative heterochromatin structure and gene repression in this fungus, including core PRC2 subunits and accessory proteins, the histone variant H2A.Z, the chromatin remodeler IMITATION SWITCH (ISW), the H3K36 methyltransferase ASH1, telomere binding proteins, and components of the constitutive heterochromatin pathway (Jamieson et al. 2013; Basenko et al. 2015; Jamieson et al. 2016; Bicocca et al. 2018; Jamieson et al. 2018; Courtney et al. 2020; McNaught et al. 2020; Kamei et al. 2021; Wiles et al. 2022; Mumford et al. 2024; Ebot-Ojong et al. 2025). In contrast to mammals, where H3K36me3 antagonizes PRC2, ASH1 catalyzed H3K36 methylation co-localizes with H3K27me3 in *N. crassa* and other fungi (Yuan et al. 2011; Schmitges et al. 2011; Bicocca et al. 2018; Janevska et al. 2018; Alabert et al. 2020; Moller et al. 2023; Xu et al. 2024). Moreover, ASH1 catalytic activity is required for gene repression in PRC2-repressed domains (Janevska et al. 2018; Bicocca et al. 2018; Xu et al. 2024). In addition, the H3K27me3 reader protein EPR1 and the RPD3L histone deacetylase complex are required for normal gene repression in facultative heterochromatin but not for normal H3K27me3 patterns (Mumford et al. 2024; Wiles et al. 2020).

Despite major advances in our understanding of facultative heterochromatin formation in filamentous fungi, the role of histone chaperones remains unexplored. Histone chaperones are a group of chromatin regulators that bind to newly synthesized or recycled histones and assemble nucleosomes on DNA (Hammond et al. 2017). Histone chaperones are generally categorized into two groups, replication-dependent or replication-independent, based on their activity inside or outside of S-phase (Tagami et al. 2004; Nakatani et al. 2004; Kollenstart et al. 2026). Proper function of replication-dependent chaperones is critical for maintenance of certain epigenetic modifications over multiple mitotic divisions (Alabert and Groth 2012; Hammond et al. 2017; Dreyer et al. 2024). Chromatin Assembly Factor 1 (CAF-1) is a highly conserved, heterotrimeric, replication-dependent histone chaperone complex (Smith and Stillman 1989). CAF-1 is recruited to replication forks and sites of DNA double-strand breaks (DSBs) through the interaction of its large subunit (called CAC-1 in *N. crassa*) with the DNA processivity factor Proliferating Cell Nuclear Antigen (PCNA) (Shibahara and Stillman 1999; Moggs et al. 2000; Zhang et al. 2000; Ransom et al. 2010). CAF-1 receives newly synthesized H3/H4 dimers from upstream histone chaperones and deposits H3/H4 tetramers onto nascent DNA (Smith and Stillman 1989, 1991; Sauer et al. 2018; Liu et al. 2023; Rouillon et al. 2023). Previous work in *S. cerevisiae* and *S. pombe* has implicated CAF-1 and other replication-dependent chaperones in gene silencing and heterochromatin maintenance (Enomoto and Berman 1998; Huang et al. 2007; Dohke et al. 2008). Studies in *Drosophila*, *Arabidopsis thaliana*, and mouse Embryonic Stem Cells (mESCs) suggest the CAF-1 complex is important for proper gene silencing, H3K27me3 maintenance, and cellular plasticity (Houlard et al. 2006; Schonrock et al. 2006; Song et al. 2007; Huang et al. 2010; Ishiuchi et al. 2015; Mozgova et al. 2015; Hatanaka et al. 2015; Cheloufi et al. 2015; Jiang and Berger 2017; Roelens et al. 2017; Cheng et al. 2019; Franklin et al. 2022; Dreyer et al. 2024). However, CAF-1 knockouts exhibit severe developmental defects in these systems and are embryonic lethal in mammals, making it difficult to study the role of CAF-1 in facultative heterochromatin formation and maintenance (Hoek and Stillman 2003; Schonrock et al. 2006; Song et al. 2007). Indeed, the mechanisms by which CAF-1 contributes to stable, PRC2-dependent gene repression remain unclear. Genetic and biochemical studies in plants and mammalian cells support a direct role for CAF-1 in promoting H3K27me3 deposition (Cheng et al. 2019; Jiang and Berger 2017). In contrast, cell cycle analyses of H3K27me3 recovery kinetics found that *de novo* establishment of H3K27me3 occurs primarily during the subsequent G1 phase rather than during DNA replication (Reverón-Gómez et al. 2018). Thus, additional work is needed to determine how CAF-1 contributes to normal H3K27me3 patterns across eukaryotes.

Like PRC2, the CAF-1 complex is fully conserved in *N. crassa*, where it is non-essential for viability. Here, we report that mutation of individual CAF-1 subunits results in derepression of H3K27me3-marked genes in *N. crassa*. This is accompanied by widespread changes to the chromatin environment, including a loss of H3K36me3 at a subset of genes within facultative heterochromatin domains and redistribution of H3K27me3 within the genome. Double mutants lacking both CAF-1 and PRC2 components displayed more severe gene silencing and chromatin defects than either single mutant, suggesting that CAF-1 and PRC2 play an overlapping, rather than a linear role in maintaining facultative heterochromatin. Together, these results identify CAF-1 as an important regulator of facultative heterochromatin and reveal a functional link between replication-coupled nucleosome assembly and PRC2-dependent gene repression in *N. crassa*.

## Results

### The CAF-1 complex is required for normal repression of genes within facultative heterochromatin

Based on previous research in other model systems, we hypothesized that one or more histone chaperones would be required for proper establishment or maintenance of facultative heterochromatin in *N. crassa* (Hammond et al. 2017). To test this, we analyzed RNA-seq data from gene deletion strains lacking individual histone chaperones available from the *N. crassa* gene deletion (Dunlap et al. 2007). We first plotted the average transcript level of 506 H3K27me3-marked genes normally repressed in wild type mycelia (H3K27me3-repressed), which revealed that mutants lacking individual subunits of the CAF-1 complex (*Δcac-1*, *Δcac-2*, *Δcac-3*) displayed significantly elevated transcript levels for this gene set (Fig. 1A; Dataset S1A). The average expression level of H3K27me3-marked genes in individual CAF-1-deficient mutants was comparable to the Δset-7 control strain (lacking the catalytic subunit of PRC2; Fig. 1A). H3K27me3-marked genes were not broadly induced in deletion strains lacking other histone chaperones (Fig. 1A; Dataset S1A). Expression profiles of replicate samples were highly similar (Figure S1A-B).

**Figure 1.**
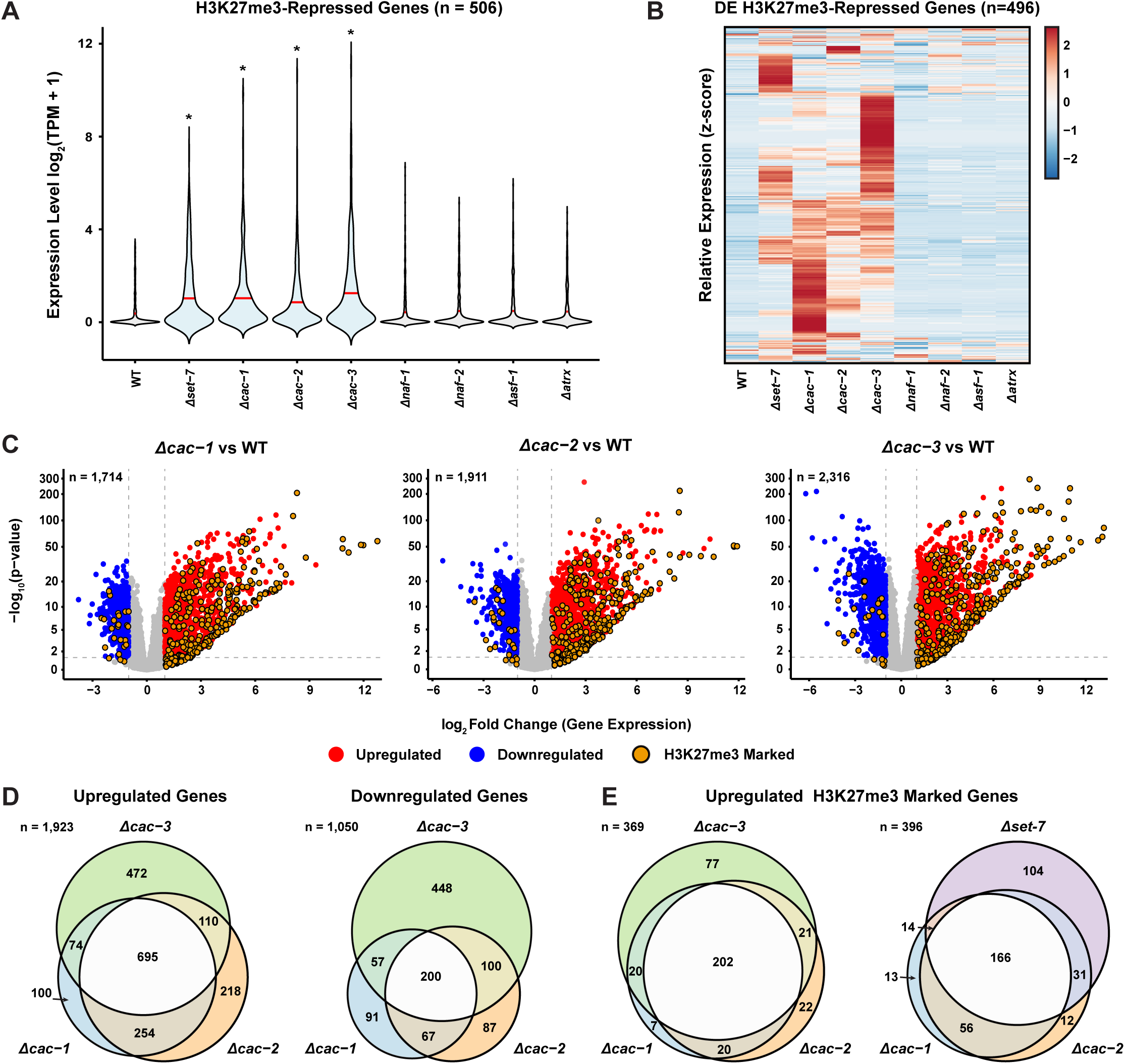
CAF-1 deficiency results in derepression of H3K27me3-marked genes. **(A)** The violin plot shows expression level of 506 H3K27me3-repressed genes in wild type and the indicated histone chaperone-deficient mutants. The “*” indicates a significant difference in average expression level relative to wild type (Wilcoxon test). **(B)** The heatmap shows relative expression levels of 496 differentially expressed, H3K27me3-repressed genes in wild type and the indicated histone chaperone-deficient mutants. Data are row-normalized and genes (rows) are hierarchically clustered (columns). **(C)** The volcano plots show relative gene expression (log_2_[mutant/wild type], x-axis) and adjusted p-value (–log_10_[adjusted p-value], y-axis) for all *N. crassa* genes in CAF-1-deficient mutants compared to wild type. “n” represents total number of differentially expressed genes. Non-differentially expressed, non-H3K27me3-marked genes are colored gray. **(D)** The Venn diagrams display overlap of genes significantly upregulated (left) or downregulated (right) in CAF-1-deficient mutants compared to wild type. “n” represents total number of genes. **(E)** The Venn diagrams display overlap of H3K27me3-marked genes significantly upregulated in CAF-1-deficient and PRC2-deficient. “n” represents total number of genes.

To examine the expression of individual H3K27me3-marked genes, we constructed a heatmap plotting relative transcript levels in wild type and mutant strains (Fig. 1B; Dataset S1.A). This revealed widespread upregulation of H3K27me3-marked genes in CAF-1-deficient mutants and Δ*set-7.* The gene expression profile in CAF-1-deficient mutants was distinct from that of *Δset-7*. In total, we found that 2,973 genes were differentially expressed across all three CAF-1-deficient mutants relative to wild type (∼28% of the genome), with H3K27me3-marked genes representing some of the most highly upregulated genes (Fig. 1C-D; Dataset S1B-E). The transcript profile was similar for *Δcac-1* and *Δcac-2*, which displayed overlap for ∼65% of genes upregulated across both mutants (949/1451 genes; Fig. 1D). This concordance likely reflects the fact that disrupting any of the CAF-1 subunits diminishes H3/H4 binding activity of the remaining components (Kaufman et al. 1995; Liu et al. 2016; Sauer et al. 2018). The transcript profile of Δ*cac-3* was distinct, likely because CAC-3 and its homologs are present in multiple chromatin modifying complexes, including the SIN3 and RPD3S/L histone deacetylase (HDAC) complexes and PRC2 (Fig. 1B,D) (Zhang et al. 1997; Martinez-Balbas et al. 1998; Tong et al. 1998; Xue et al. 1998; Jamieson et al. 2013; Tang et al. 2021; Lin et al. 2022). Overall, all three CAF-1-deficent mutants displayed substantial overlap of upregulated H3K27me3-marked genes, with 202 of the 369 genes upregulated in all three strains (Fig. 1E). CAF-1 mutants also displayed upregulation of genes within euchromatin regions. Gene Ontology (GO) analysis of genes that were differentially regulated in CAF-1-deficient strains did not link CAF-1 to any specific biological processes, though some upregulated categories are associated with fruiting body development (Fig. S1C; “ascospore release from ascus”), consistent with previously reported findings in a PRC2-deficient strain (Deaven et al. 2025). Taken together, these data suggest that CAF-1-deficiency results in widespread misregulation of gene expression, including within facultative heterochromatin domains.

### CAF-1 is required for normal patterns of repressive histone methylation in facultative heterochromatin

To determine if increased gene expression in facultative heterochromatin is associated with altered histone modification profiles, we performed ChIP-seq to examine H3K27me3 enrichment in individual CAF-1-deficient mutants and a wild-type control strain. Using the IGV genome browser, we visualized H3K27me3-enrichment patterns and the genome locations of mis-expressed genes in each mutant strain (Fig. 2A). In wild type, facultative heterochromatin domains are large, multi-gene domains enriched for H3K27me3. Within these regions, clusters of adjacent genes were upregulated in CAF-1-deficient mutants. These mutants also displayed reduced H3K27me3 enrichment at a subset of facultative heterochromatin regions (Fig. 2A-B). Within euchromatin, misregulated genes were not clustered at specific chromosomal regions (Fig 2A; Dataset S1.B-D). To visualize the genome-wide patterns of H3K27me3, we quantified H3K27me3 enrichment in 300 base-pair windows and plotted the level of enrichment for the individual CAF-1-deficient strains and the wild-type and Δ*set-7* control strain. This revealed that CAF-1-deficient mutants exhibit regional losses of H3K27me3 as well as ectopic gains at novel loci (Fig. 2C; Dataset S2A). The *Δcac-3* strain displayed a sub-telomeric loss and internal ectopic gains of H3K27me3 as previously reported, and the patterns of H3K27me3 enrichment in the *Δcac-1* and *Δcac-2* strains were distinct from *Δcac-3*, consistent with their gene expression profiles (Fig. 2A-C) (Jamieson et al. 2013; Courtney et al. 2020). The *Δcac-1* and *Δcac-2* profiles were highly similar, suggesting that disruption of CAF-1 complex is responsible for the H3K27me3 defect observed in these strains. Independent validation of select regions by ChIP-qPCR supported ChIP-seq observations, and replicate experiments using an alternative H3K27me3 antibody produced identical results (Fig. S2). Measuring H3K27me3 levels by western blot revealed that CAF-1 deficiency did not result in a global decrease of H3K27me3, indicating that aberrant gains of H3K27me3 offset regional losses (Fig. 2D). Notably, we did not observe altered expression for any genes encoding known components of the facultative heterochromatin pathways (Dataset S1.B-D).

**Figure 2.**
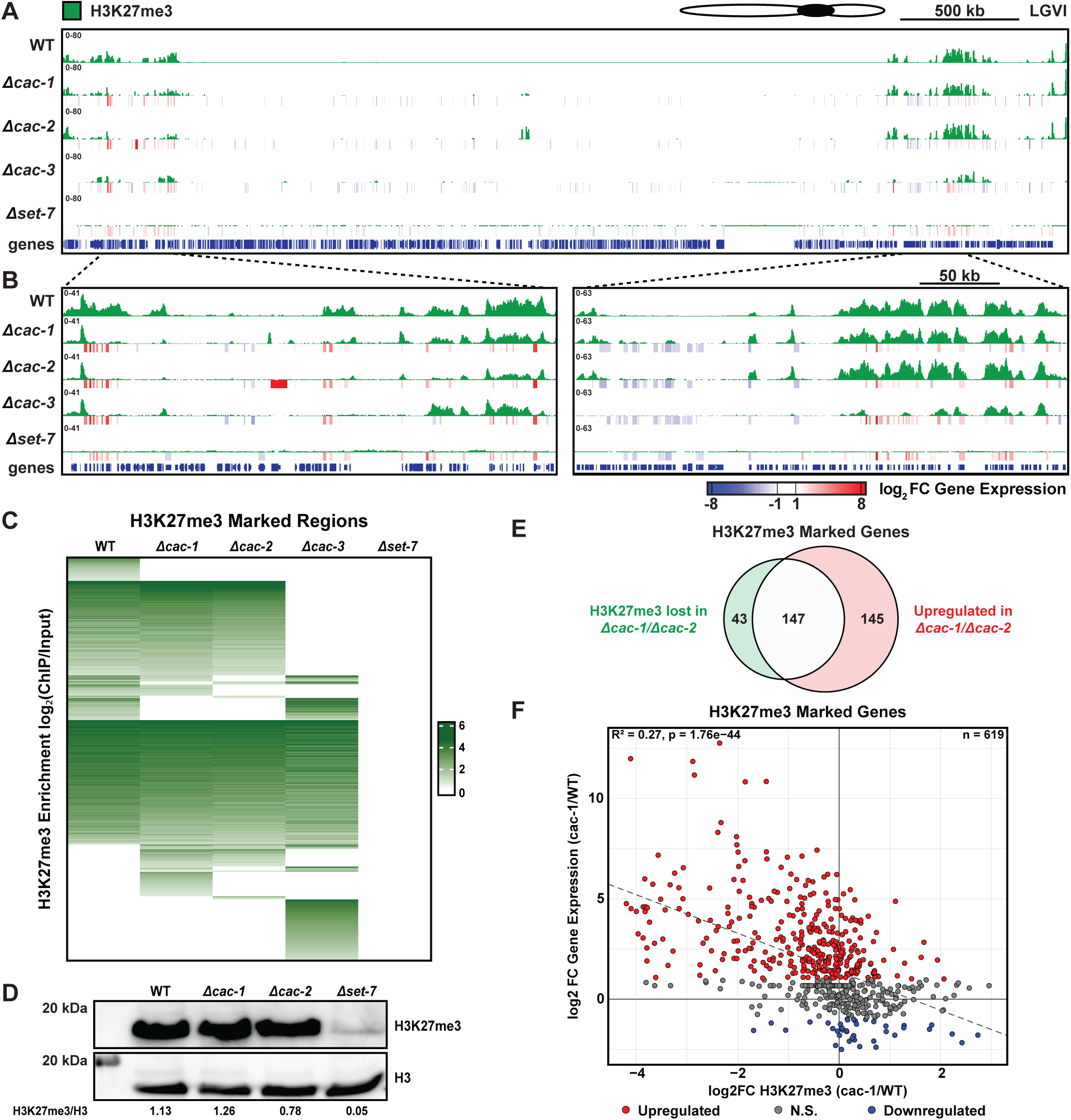
CAF-1 deficiency results in aberrant H3K27me3 patterns. **(A)** A whole-chromosome view of H3K27me3 enrichment and relative gene expression (log_2_[mutant/wild type]) is shown for linkage group VI. **(B)** Magnified views of H3K27me3 enrichment and relative gene expression (log_2_[mutant/wild type]) are shown for two facultative heterochromatin domains on linkage group VI. **(C)** The heatmap displays H3K27me3 enrichment (log₂[ChIP/Input]) in wild type, CAF-1 deficient mutants, and *Δset-7* across H3K27me3-marked regions. Values with an FDR > 0.05 were set to 0. Rows correspond to 300 bp bins within H3K27me3-marked loci and are manually clustered and sorted by enrichment in wild type and mutant strains. **(D)** Western blots were performed using whole-cell extract and probed with antibodies against H3K27me3 or total histone H3 for the indicated strains. Numbers represent H3K27me3 band intensities quantified by densitometry and normalized to total H3 band intensity. **(E)** The Venn diagram shows overlap of H3K27me3-marked genes that lose H3K37me3 with genes upregulated in *Δcac-1* or *Δcac-2*. **(F)** The scatter plot shows relative H3K27me3 enrichment (log_2_[Δ*cac-1*/wild type], x-axis) compared to relative gene expression (log_2_[Δ*cac-1*/wild type], y-axis) for H3K27me3-marked genes.

We next asked whether H3K27me3 losses were correlated with increased transcription of H3K27me3-marked genes. For 302 genes that reside in facultative heterochromatin and were upregulated in *Δcac-1* or *Δcac-2*, we found that ∼50% showed reduced H3K27me3 levels in their promoters (Fig. 2E; Datasets S1, S2B). We also identified 43 genes that lost H3K27me3 in CAF-1-deficient mutants but were not induced. We next plotted differential enrichment of H3K27me3 and compared this to differential gene expression data for 619 protein coding genes enriched for H3K27me3 in the wild type and/or Δ*cac-1* strain. We observed a weak correlation between loss of H3K27me3 and upregulated gene expression (Fig. 2F, Fig. S3A-B). Additionally, 74 protein-coding genes were located at sites of ectopically gained H3K27me3 in *Δcac-1*. Only 13 were significantly downregulated. Thus, loss of H3K27me3 is not solely responsible for increased expression of facultative heterochromatin genes, and conversely, loss of H3K27me3 is not sufficient to drive increased expression in CAF-1-deficient mutants. Together, these data suggest that CAF-1 mutants have an altered facultative heterochromatin environment, but that perturbing the complex does not result in a global loss of H3K27me3. The Δ*cac-3* strains exhibited a distinct phenotype compared to Δ*cac-1* and Δ*cac-2,* presumably because CAC-3 is a component of multiple complexes, including PRC2. We therefore decided to focus on *Δcac-1* and *Δcac-2* to specifically probe the role of CAF-1 in facultative heterochromatin.

We next asked if CAF-1 might be required for proper patterning of other repressive histone modifications. In wild type, H3K36me3 is broadly enriched in facultative heterochromatin covering both promoters and gene bodies (Bicocca et al. 2018; Kamei et al. 2021). ChIP-seq for H3K36me3 revealed that CAF-1-deficient mutants display regional losses of H3K36me3 in facultative heterochromatin (Fig. 3A). On average, H3K36me3 levels across facultative heterochromatin genes were lower in Δ*cac-1* and Δ*cac-2* than in wild-type or *Δset-7* strains (Fig. 3B). In contrast, the Δ*cac-1* and Δ*cac-2* strains had higher H3K36me3 levels in the coding sequence of euchromatic genes targeted by SET-2 (Adhvaryu et al. 2005) (Fig. 3B). Differential enrichment analysis identified 76 gene promoters with significantly reduced H3K36me3 in *Δcac-1* and Δ*cac-2* (Fig. 3C, “Lost in CAF-1 category”; Dataset S2C). Most of these genes retained gene body H3K36me3, consistent with co-transcriptional deposition of H3K36me3 by SET-2. The *Δset-7* mutant displayed a loss of H3K36me3 at only 33 of these 76 gene promoters (Fig. 3D; Dataset S2C). We found that for 63/76 genes, expression increased in *Δcac-1* or *Δcac-2* (Fig. 3D, Datasets S1, S2C). However, plotting relative H3K36me3 enrichment against relative gene expression in CAF-1-deficient mutants revealed a weaker correlation than that observed for H3K27me3 enrichment, while *Δset-7* displayed no obvious correlation (Fig. S4). Reduced H3K36me3 levels in promoters frequently co-occurred with reduced H3K27me3-enrichment (51/76 genes, Fisher’s exact test, odds ratio=4.80, p<4.97x10^-10^; Fig 3D; S5A-B). Furthermore, 256 of the 292 H3K27me3-marked genes upregulated in *Δcac-1* or *Δcac-2* are also upregulated in *ash-1(Y888F)* (87%; Fisher’s exact test, odds ratio=4.31, p<1.79x10^-12^ Fig. 3D, Datasets S1, S2D). (Bicocca et al. 2018). This represents a higher percent overlap than that observed for *Δcac-1* or *Δcac-2* and *Δset-7* (∼67%), or for *Δset-7* and *ash1(Y888F)* (∼74%) (Fig. 1E, S5D).

**Figure 3.**
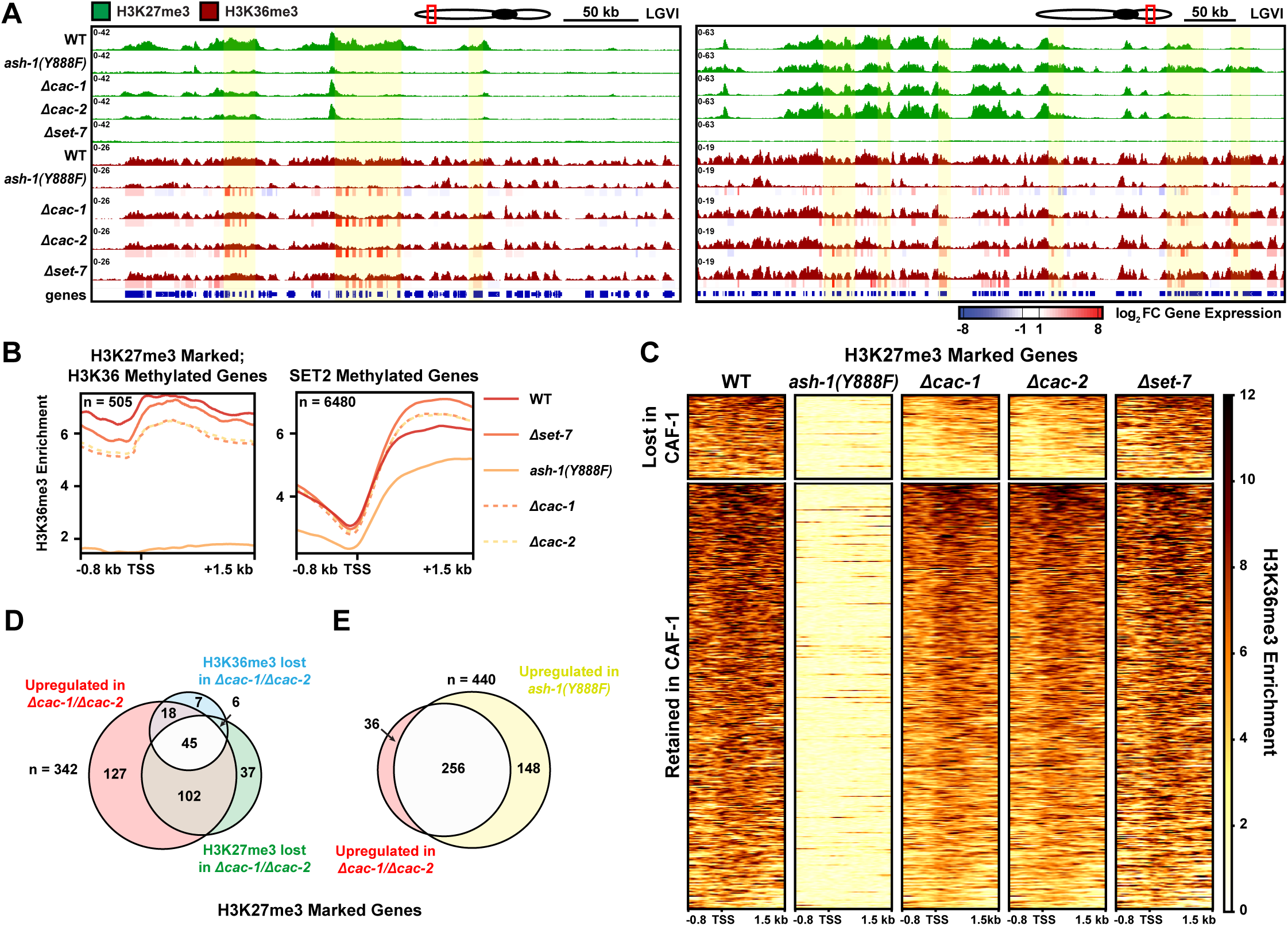
CAF-1 deficiency perturbs ASH1-dependent H3K36me3. **(A)** The genome browser images show enrichment of H3K27me3, H3K36me3, and relative gene expression (log_2_[mutant/wild type]) at two facultative heterochromatin domains on linkage group VI. The *ash-1(Y888F)* data track shows published data for H3K27me2/3 enrichment. The highlighted regions exhibit H3K36me3 loss in one or more mutants. **(B)** The metaplots show mean H3K36me3 enrichment from −0.8 kb to +1.5 kb relative to the TSS for two gene sets: H3K27me3-marked and SET-2–methylated genes. “n” = total number of genes in each category. CAF-1-deficient mutants (*Δcac-1*, *Δcac-2*) are represented by dashed lines and controls (wild type, *Δset-7*, *ash-1(Y888F)*) are represented by solid lines. **(C)** The heatmaps show H3K36me3 enrichment from −0.8 kb to +1.5 kb relative to the TSS for H3K36me3-enriched, H3K27me3-marked genes (n = 505). Clusters represent genes that lose (top) or retain (bottom) H3K36me3 in *Δcac-1* and *Δcac-2*. Clusters are individually sorted by the sum of H3K36me3 enrichment level across all strains. **(D)** The Venn diagram shows the number of genes that lose H3K36me3, lose H3K27me3, or are upregulated in *Δcac-1* and *Δcac-2* (left) or (**E**) genes that are upregulated in *Δcac-1* and *Δcac-2* or upregulated in *ash-1(Y888F)*. “n” represents total number of genes.

Other repressive modifications were not affected in the CAF-1-deficient strains. ChIP-seq analysis of H4K20me3 revealed that the PTM is enriched over most of the genome but excluded from H3K9me3-enriched constitutive heterochromatin domains. No obvious changes in H4K20me3 patterns were observed in *Δcac-1*, *Δcac-2*, or *Δset-7* compared to wild type (Fig. S6). Previous work in other organisms found that CAF-1-deficiency results in altered H3K9me3 at constitutive heterochromatin and revealed a direct interaction between CAF-1 and Heterochromatin Protein 1 (HP1), a H3K9me3 reader protein (Murzina et al. 1999; Thiru et al. 2004; Houlard et al. 2006; Dohke et al. 2008; Cheloufi et al. 2015). ChIP-seq experiments to examine H3K9me3 patterns in CAF-1-deficient mutants revealed no significant changes, suggesting CAF-1 is not required for normal constitutive heterochromatin in *N. crassa* (Fig. S6). Taken together, these data show that CAF-1 is required for normal repressive histone methylation patterns in facultative heterochromatin, but not within constitutive heterochromatin domains.

### CAF-1 deficiency leads to gain of active PTMs within facultative heterochromatin regions

Considering the transcriptional misregulation and defects in facultative heterochromatin structure observed in the Δ*cac-1* and -*2* strains, we next asked if CAF-1 deficiency leads to increased histone PTMs associated with active transcription at H3K27me3-marked genes. We decided to examine H3K4me2 and pan-lysine acetylation (Kac), PTMs known to antagonize PRC2 activity (Tie et al. 2009; Creyghton et al. 2010; Pasini et al. 2010; Schmitges et al. 2011). Within typically repressed, H3K27me3-marked genes, Kac was gained in a subset of facultative heterochromatin genes. A total of 104 genes exhibited elevated promoter Kac levels in *Δcac-1* or *Δcac-2* (Fig. 4A-D; Dataset S3A). Kac patterns were distinct between CAF-1- and PRC2-deficient mutants, with only ∼11.5% of genes with increased Kac in *Δcac-1* or *Δcac-2* exhibiting gains in *Δset-7*, which also displayed elevated Kac at 37 unique genes (Fig. 4D). ChIP-seq analysis of H3K4me2 revealed gains in 24 H3K27me3-marked genes in *Δcac-1* or *Δcac-2* (Fig. 4A-D, Fig. S7A; Dataset S3B). The Δ*set-7* strain gained H3K4me2 in 22 of these 24 genes as well as 26 other H3K27me3-marked genes (Fig. 4D, Fig. S7A). The *Δcac-1* and *Δcac-2* mutants displayed significant overlap in enrichment of both H3K4me2 and Kac, with 60 of the 62 genes that gain either modification in *Δcac-1* also exhibiting gains in *Δcac-2*, suggesting that exclusion of these active modifications from facultative heterochromatin depends on the CAF-1 complex (Fig. S7). Surprisingly, *Δcac-2* displayed gains of H3K4me2 or Kac at an additional 60 genes compared to *Δcac-1* (Fig. S7C). Gene expression of H3K27me-enriched genes displayed a weak correlation with enrichment of Kac in both *Δcac-1* and *Δcac-2* (Fig. 4E, S7E). Moreover, plotting relative H3K27me3 or H3K36me3 enrichment against relative Kac enrichment in CAF-1-deficient mutants (log_2_[mutant/wild type]) revealed weak negative correlations (Fig.4 F-G, Fig.S7F-G). Importantly, complementing the *Δcac-1* and Δ*cac-2* mutations restored wild type chromatin modification profiles at facultative heterochromatin domains (Fig. S8). Overall, the gains in active modifications are modest compared to the losses of H3K27me3 and H3K36me3 in gene promoters. Collectively, these data suggest that upregulation of gene expression within facultative heterochromatin domains of CAF-1-deficient mutants is primarily due to losses of repressive modifications, but that a subset of expressed genes gain PTMs typically associated with active transcription.

**Figure 4.**
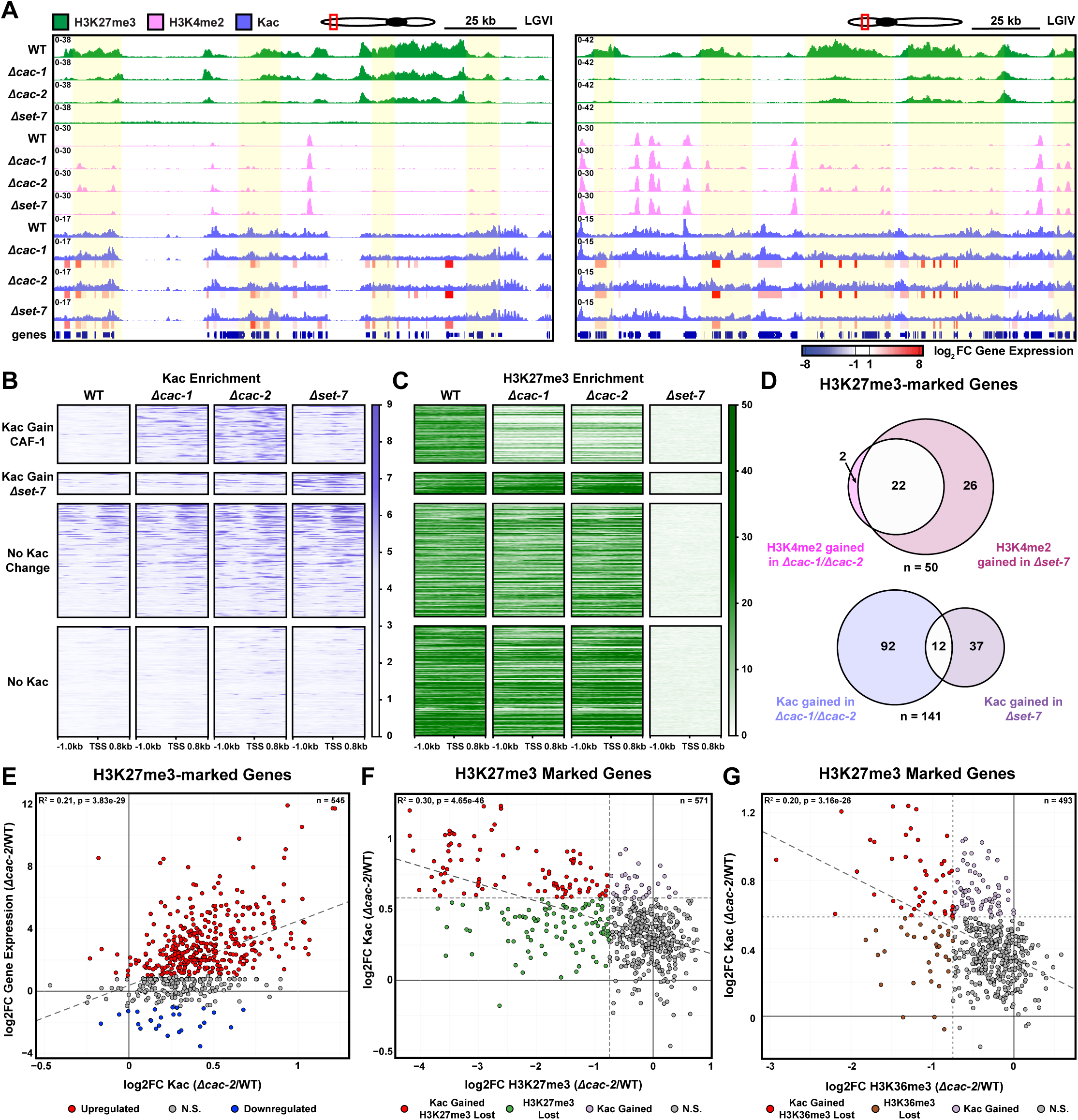
CAF-1 deficiency results in an euchromatin-like environment at H3K27me3-marked genes. **(A)** The zoomed-in browser shots show H3K27me3, H3K4me2, Kac, and relative gene expression (log_2_[mutant/wild type]) for the indicated strains at two facultative heterochromatin domains on linkage groups IV and VI. Highlights represent regions of H3K4me2 or Kac gain in one or more mutants. **(B-C)** The heatmaps display: Kac (B) or H3K27me3 (C) from −1 kb to +0.8 kb relative to the TSS at typically H3K27me3-marked genes (n= 571). Genes (rows) are split into four clusters (top to bottom): Kac gained in CAF-1-deficient mutants (1), Kac gained in *Δset-7* but not in CAF-1-deficient mutants (2), no change in Kac vs wild type (3), and unacetylated in all strains (4). **(D)** Venn diagrams displaying overlap in H3K27me3-marked genes that gain H3K4me2 (top) or gain Kac (bottom) in Δcac-1/Δcac-2 or *Δset-7*. “n” = total number of genes. **(E)** The scatter plot shows relative Kac enrichment (log_2_[Δ*cac-2*/wild type], x-axis) against relative gene expression (log_2_[Δ*cac-2*/wild type], y-axis) for typically H3K27me3-marked genes. (**F)** The scatter plot shows relative H3K27me3 enrichment (log_2_[Δ*cac-2*/wild type], x-axis) against relative Kac enrichment (log_2_[Δcac-2/wild type], y-axis) for typically H3K27me3-marked genes. (**G**) The scatter plot shows relative H3K36me3 enrichment (log_2_[Δ*cac-2*/wild type], x-axis) against relative Kac enrichment (log_2_[Δcac-2/wild type], y-axis) for typically H3K27me3-marked genes. “n” = total number of genes.

### CAF-1 and PRC2 play distinct roles in gene repression

In *Arabidopsis thaliana* and mESCs, CAF-1 is proposed to interact directly with PRC2 to establish or maintain normal H3K27me3 patterns (Jiang and Berger 2017; Cheng et al. 2019). However, we found that Δ*set-7* and CAF-1-deficient strains show upregulation of different sets of genes within facultative domains, suggesting that PRC2 and CAF-1 function separately to repress genes within facultative heterochromatin (Fig. 1B, E). To test this possibility, we generated a double mutant lacking *cac-1* and the PRC2 subunit *suppressor of zeste-12* (*Δcac-1; Δsuz-12*), and we performed RNA-seq analysis to examine expression of genes residing in facultative heterochromatin domains (Dataset S4). The *Δcac-1; Δsuz-12* strain exhibited a synergistic silencing defect, with 473 of the 506 H3K27me3-marked genes (∼93%) exhibiting significant upregulation compared to wild type (Fig. 5). Notably, *Δcac-1; Δsuz-12* displayed upregulation of nearly every H3K27me3-marked gene upregulated in *Δcac-1* or *Δsuz-12*, in addition to 146 genes that were not upregulated in either single mutant, suggesting that CAF-1 and PRC2 do not function in a linear pathway to repress genes within *N. crassa* facultative heterochromatin regions. Replicate RNA-seq samples produced highly similar data (Fig. S9). Interestingly, 4,577 genes were differentially expressed in *Δcac-1; Δsuz-12* (∼47% of the genome), 2,699 of which were upregulated, suggesting that combined CAF-1 and PRC2-deficiency results in a genome-wide misregulation of gene expression (Fig. 5D). GO analysis revealed that genes misregulated in *Δcac-1; Δsuz-12* are involved in diverse biological processes such as DNA replication and repair, translation, and stress response (Fig. S9).

**Figure 5.**
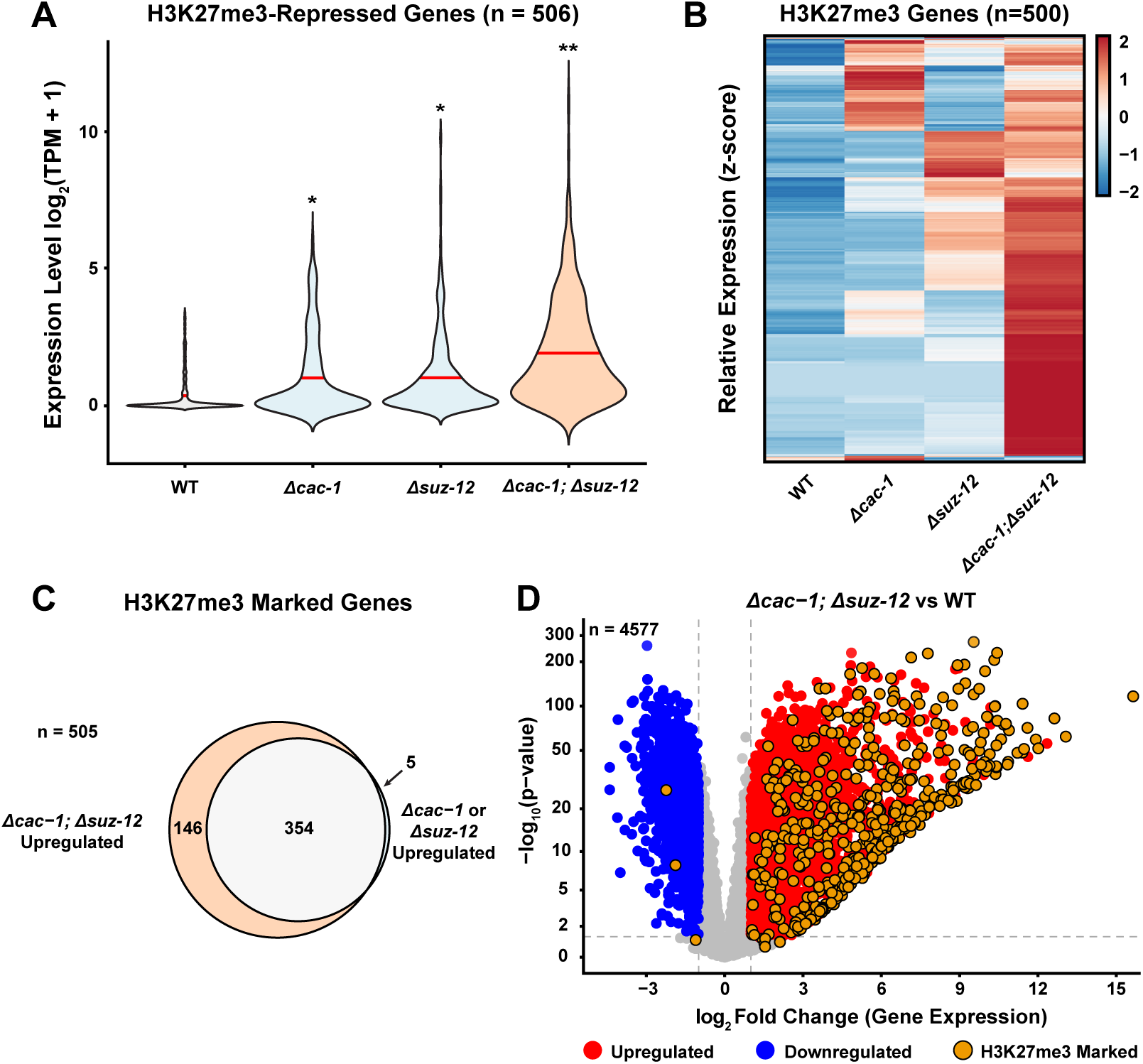
CAF-1 and PRC2 play distinct roles in H3K27me3-marked gene repression. **(A)** The violin plot shows expression level of 506 H3K27me3-repressed genes in *Δcac-1*, *Δsuz-12*, and *Δcac-1; Δsuz-12*. * and ** indicates significant difference in average expression level across all plotted genes compared to wild type based on Wilcoxon test (* = p-value < 1.5 x 10^-14^, ** = p-value < 1.04 x 10^-88^). **(B)** The heatmap shows relative expression level of 500 H3K27me3-repressed genes that are differentially expressed in wild type, *Δcac-1*, *Δsuz-12*, or *Δcac-1; Δsuz-12*. Data are row-normalized, and genes (rows) are hierarchically clustered (columns). **(C)** The Venn diagram shows overlap of H3K27me3-marked genes upregulated in *Δcac-1* or *Δsuz-12* and *Δcac-1; Δsuz-12*. “n” represents total number of genes. **(D)** The volcano plot shows relative gene expression (log_2_[Δ*cac-1; Δsuz12*/wild type], x-axis) plotted against adjusted p-value (-log_10_[adjusted p-value], y-axis) for all *N. crassa* genes. “n” represents total number of differentially expressed genes. Non-differentially expressed, non-H3K27me3-marked genes are colored gray.

We next asked if global defects in gene expression resulted in phenotypic changes in *Δcac-1; Δsuz-12*. The growth rate of the Δ*suz-12* and Δ*cac-1* strains was similar to wild type, whereas the *Δcac-1; Δsuz-12* strain displayed a significant growth defect on minimal medium (Fig. S10). In contrast, the DNA-damage sensitivity of the Δ*cac-1; Δsuz-12* was similar to the *Δcac-1* single mutants (Fig. S10C). Collectively, these data suggest that CAF-1 and PRC2 play distinct but complementary roles in maintaining gene repression of H3K27me3-marked genes.

## Discussion

Re-establishment of histone PTMs following DNA replication is essential for maintenance of repressive chromatin states and long-term epigenome stability (Kollenstart et al. 2026). Accordingly, how histone chaperones contribute to faithful inheritance of chromatin states has been a subject of intense investigation. Using the genetically tractable fungus *Neurospora crassa*, we show that the replication-dependent histone chaperone CAF-1 is required for normal patterns of repressive histone PTMs and for gene repression in facultative heterochromatin.

Genetic and biochemical studies in plants and mammalian cells have suggested that CAF-1 physically associates with PRC2 to facilitate H3K27me3 deposition (Cheng et al. 2019; Jiang and Berger 2017). Given that loss of CAF-1 disrupts facultative heterochromatin in animals, plants, and *N. crassa*, it is tempting to speculate that CAF-1 regulates PRC2-repressed chromatin through conserved mechanisms. However, several observations argue against a simple model in which CAF-1 maintains H3K27me3 solely through direct, positive regulation of PRC2. Here, we found that simultaneous loss of CAF-1 and PRC2 caused a synergistic silencing defect in *N. crassa*, suggesting that these complexes function through partially independent mechanisms. Furthermore, Reverón-Gómez and colleagues showed that re-establishment of H3K27me3 in mammalian cells occurs primarily during G1 phase, rather than during S phase when CAF-1 activity is highest (Reverón-Gómez et al. 2018). Together, the available data suggest that CAF-1 contributes to facultative heterochromatin maintenance through broader effects on chromatin organization and not only through direct regulation of PRC2.

CAF-1 plays a central role in replication-coupled nucleosome assembly (Hammond et al. 2017; Liu et al. 2023; Kollenstart et al. 2026). We propose that loss of CAF-1 delays nucleosome assembly behind the replication fork, which in turn destabilizes facultative heterochromatin in *N. crassa*. Supporting this model, CAF-1-deficient cells exhibit reduced nucleosome occupancy and delayed chromatin maturation during S phase (Fennessy and Owen-Hughes 2016; Munoz-Viana et al.

2017; Barrientos-Moreno et al. 2023; Chen et al. 2023). This reduced nucleosome occupancy may impair the catalytic activity of ASH1, PRC2, or both. Indeed, mammalian PRC2 activity is stimulated by higher nucleosome density, and in *N. crassa*, ASH1-dependent H3K36 methylation is lost in the absence of ISW, a nucleosome sliding chromatin remodeler (Kamei et al. 2021; Yuan et al. 2012).

Alternatively, delayed nucleosome assembly during S phase may create an extended permissive chromatin state in which transcription factors, RNA polymerase II, and other transcriptional machinery gain access to promoters before repressive chromatin is fully restored (Alvarez et al. 2023). The observation that CAF-1 mutants displayed widespread transcriptional upregulation within both facultative heterochromatin and euchromatin is consistent with this idea. Replication-independent histone chaperones are thought to compensate for loss of CAF-1 activity, and studies in HeLa cells showed that reduced CAF-1 activity promotes HIRA-dependent incorporation of histone H3.3 at sites enriched for RNA polymerase II (Ray-Gallet et al. 2011; Barrientos-Moreno et al. 2023). Although our data do not distinguish whether the loss of repressive histone PTMs precedes transcriptional activation or instead results from aberrant transcription, our results are consistent with a model in which defective replication-coupled nucleosome assembly destabilizes facultative heterochromatin and promotes inappropriate transcription.

In addition to widespread transcriptional misregulation, we found that CAF-1-deficient strains exhibit reduced ASH1-dependent H3K36 methylation and extensive redistribution of H3K27me3. The mechanisms underlying H3K27me3 redistribution remain unclear. H3K27me3 redistribution may occur because PRC2 complexes are displaced from transcriptionally activated loci, leading to off-target methylation at other regions of the genome. Indeed, ectopic H3K27me3 has been observed when other chromatin regulators are disrupted (Basenko et al. 2015; Bicocca et al. 2018; Kamei et al. 2021; Lee et al. 2023; Ebot-Ojong et al. 2025). These observations are consistent with a “source-sink” model in which PRC2 is normally concentrated at a defined set of facultative heterochromatin domains (“sources”) but redistributes to ectopic loci (“sinks”) when chromatin repression is perturbed (Murphy and Berger 2023). Taken together, our findings show that CAF-1 is required for gene repression within facultative heterochromatin and support a model in which replication-coupled nucleosome assembly is required to preserve stable PRC2-repressed chromatin states.

## Materials and Methods

### Strain generation and growth conditions

All strains used in this study are listed in Table S1. Oligos used for strain generation, and genotyping are listed in Table S2. Double mutants were generated through crossing of single mutants as previously described (Davis and de Serres 1970). Complementation strains were generated by transforming full length, 3xFLAG-tagged gene constructs into the his-3 locus of strains harboring a deletion allele at the native locus. The *his-3^+^::cac-1-3xflag*; Δ*cac-1* complemented strain was generated by introducing the *cac-1-3xflag* construct into the *his-3* locus of an otherwise wild type strain and then crossing the resulting transformant to Δ*cac-1* because we were unable to successfully transform a *his-3;* Δ*cac-1* strain presumably due to defects in homologous recombination and DNA repair (Linger and Tyler 2005; Park et al. 2011; Gaillard et al. 1996; Green and Almouzni 2003). 3xFLAG-tagged constructs were generated via Gibson cloning using oligos in Table S2. DNA transformations were performed as previously described (Margolin et al. 1997). Strains for ChIP-seq and RNA-seq experiments were grown overnight in Vogel’s minimal media (VMM) as previously described (Davis and de Serres 1970).

### RNA isolation, ChIP-seq, and ChIP-qPCR

RNA-seq data was generated as part of a Joint Genome Institute community sequencing project (Zhou et al. 2018) or was generated in the Lewis Lab for this study. For new RNA-seq data, RNA extraction was performed as previously described (Zhou et al. 2018). Briefly, total RNA was extracted using TRIzol reagent (Thermo Fisher; Cat #:15596026) and then purified with 7.5 M LiCl. RNA purity was analyzed on agarose gels prior to library preparation. ChIP experiments were performed as previously described (Ferraro and Lewis 2018; Seymour et al. 2016), using antibodies in Table S3. ChIP samples were analyzed by sequencing on an Illumina NextSeq Novoseq instrument (Admera Health, Inc), and by qPCR (SYBR Green Universal Master Mix, Bio-Rad, 4309155). ChIP-seq libraries were constructed as previously described (Ferraro and Lewis 2018; Courtney et al. 2020). RNA-seq libraries were prepared using a NEBNext Ultra II Directional RNA Library Prep Kit according to manufacturer specifications (New England Biolabs, Catalog #E7760S).

### Protein extraction & western blotting

Cultures were grown in a shaking incubator overnight (∼18-22 hours) in liquid VMM + 1.5% sucrose at 32 °C and 180 rpm. Cultures were harvested and washed with 1x PBS, and protein was extracted by sonicating tissue on ice for 1 min. (60 sec. on, 60 sec. off; 30 µm Amplitude) in protein extraction buffer (50 mM HEPES pH 7.5, 150 mM NaCl, 1 mM EDTA, 0.02% IGEPAL, 1 mM PMSF, 1x Protease Inhibitor Cocktail; Cat #: HY-K0011). Cell lysates were centrifuged at 17,000 x *g* (∼13,200 rpm) at 4 °C, and the supernatant containing protein extract was mixed 3:1 with 4x Laemmli sample buffer (Bio-Rad, 1610747), boiled at 95 °C for 3 minutes, and run on SDS-PAGE gels at 185 V for ∼45 minutes. Western blots for 3xFLAG-tagged proteins were transferred to 0.45 µm PVDF membranes (1.5 hours wet transfer in Towbin’s buffer) and blocked for 1 hour in 5% milk in TBST, then incubated at 4°C overnight with an anti-M2-FLAG antibody (Sigma-Aldrich, F1804) in 5% milk in TBST. Western blots probing for H3K27me3 constructs were transferred to 0.45 µm PVDF membranes (7-minute Trans-Blot Turbo Transfer, 1704150) and blocked for 1 hour in 5% BSA in TBST, then incubated at 4°C overnight with an anti-H3K27me3 antibody (Diagenode, NBP2-59295) in 5% BSA in TBST. Anti-mouse and anti-rabbit primary antibodies were probed with Goat anti-Mouse IgG (H+L), HRP (ThermoFisher, #31430) and Goat anti-Rabbit IgG (H+L), HRP (ThermoFisher, 31460) secondary antibodies respectively. Membranes were developed using West-Femto Maximum Sensitivity Substrate (ThermoFisher, 34094) for 5 minutes and imaged on a Bio-Rad Chemidoc MP Imaging System.

### Bioinformatic analyses

For ChIP-seq analysis, reads were trimmed using TrimGalore! (v0.6.10) and mapped to *N. crassa* assembly 12 genome (Genbank GCA_000182925.2) using BWA (v0.7.18) “mem” (option -M) (Krueger 2012; Li and Durbin 2009). Reads were sorted and indexed using SAMtools (v1.21) (Li 2011). Bigwig files were made using deepTools (v3.5.5) “bamCoverage”, normalized by bins per million mapped reads (BPM) (Ramirez et al. 2016). Bigwig files were visualized using Integrative Genome Viewer (IGV) (Thorvaldsdottir et al. 2013) and heatmaps created using deepTools “computeMatrix” and “plotHeatmap” or the ComplexHeatmap package in R (v2.26.0) (Gu et al. 2016). Peaks were called for ChIP-seq samples using MACS3 (v3.0.1) “callpeak” (broad setting, 0.01 cutoff, 650 bp minimum peak length, 375 bp max peak gap) (Gaspar 2018). Peak used calls to determine H3K27me3-marked genes can be found in Dataset S5.

ChIP-seq enrichment was quantified using the csaw package in R (v1.42.0) and normalizing to input samples (Lun and Smyth 2016). To visualize changes to the pattern of H3K27me3 across the genome, we quantified relative enrichment in 300bp windows. H3K27me3 enrichment was determined for each gene by quantifying relative enrichment in windows that span 500 bp upstream to 300 bp downstream of each TSS. H3K36me3 enrichment was calculated from 500 bp upstream to 300 bp downstream of the TSS for H3K27me3-marked genes. Kac was calculated from 1000 bp upstream to 800 bp downstream of the TSS. H3K4me2 and enrichment was calculated over the whole gene body (TSS to TES). Genes within wild type H3K27me3 peak calls (were considered H3K27me3-enriched if they exhibited > 2 log_2_[sample/input] (4-fold change). In CAF-1-deficient strains, regions that gained H3K27me3 exhibited an enrichment value > 1 log_2_[sample/input] (2-fold change; Dataset S5). Genes were considered H3K36me3, H3K4me2, or Kac-enriched if they exhibited > 1-fold change over input. Genes were considered differently enriched if they exhibited ± 0.75 log_2_[(mutant/input)/(wild type/input)] (∼1.7-fold change) for H3K27me3, H3K36me3 and H3K4me2, and ± 0.585 log_2_[(mutant/input)/(wild type/input)] (∼1.5-fold change) enrichment for Kac.

For RNA-seq analysis, reads were trimmed with TrimGalore! and mapped using STAR (v2.7.11) “alignReads” (Dobin et al. 2013). Output files were indexed using SAMtools, and alignment counts were determined using Subread (v2.0.6) “featureCounts” (Liao et al. 2014). Differential expression was determined using the DEseq package in R (v1.48.2) (Love et al. 2014). Genes were considered differentially expressed if they exhibited ± 1 log_2_[mutant/wild type] (2-fold change) in expression compared to wild type and an adjusted p-value of < 0.05. Genes were considered “repressed” if they exhibited < 10 TPM on average across wild type samples.

### Data availability

New sequence data generated for this study is available at the National Center for Biotechnology Information’s Gene Expression Omnibus database (https://www.ncbi.nlm.nih.gov/geo) under accession numbers GSE314682 (RNA-seq) and GSE314683 (ChIP-seq). For previously published data, all Sequence Read Archive (https://www.ncbi.nlm.nih.gov/sra) accession numbers and associated strain information for samples generated at the Joint Genome Institute and other previously published datasets analyzed in this study are included in Table S4. All scripts used are publicly available on GitHub at https://github.com/eddietorres24/Torres_et_al_2026_CAF-1_Facultative_Heterochromtin_Neurospora. Supplemental dataset files have been deposited to the figshare database.

## Acknowledgements

This work was supported by the NIH/NIGMS Grant #R35GM152134 to ZAL. The work (proposal: 10.46936/10.25585/60001136) conducted by the U.S. Department of Energy Joint Genome Institute (https://ror.org/04xm1d337), a DOE Office of Science User Facility, is supported by the Office of Science of the U.S. Department of Energy operated under Contract No. DE-AC02-05CH11231.

